# ArcDFI: Attention Regularization guided by CYP450 Interactions for Predicting Drug-Food Interactions

**DOI:** 10.1101/2025.04.09.647981

**Authors:** Mogan Gim, Jaewoo Kang, Donghyeon Park, Minji Jeon

## Abstract

CYP450 isoenzymes are known to be deeply involved in the formation of drug-food interactions (DFI). Previously introduced computational approaches for predicting DFIs do not take drug-CYP450 interactions (DCI) into account and have limited generalizability in handling compounds unseen during model training. We introduce ArcDFI, a model that utilizes attention regularization guided by CYP450 interactions to predict drug-food interactions. Experimental results demonstrate ArcDFI’s ability to predict DFIs when given unseen food or drug compounds as input. Analysis of its attention mechanism provides insight into its current understanding of DCI and how they are related to its DFI predictions. To the best of our knowledge, ArcDFI is the first DFI prediction model that incorporates the concept of DCI, resulting in improved predictive generalizability and model explainability. ArcDFI is available at https://github.com/KU-MedAI/ArcDFI

## Introduction

Individuals commonly use medications to maintain health or treat diseases, but drug efficacy can be significantly influenced by dietary factors. Food components may alter drug absorption, metabolism, and excretion, potentially causing unexpected side effects, known as drug-food interactions (DFIs). DFIs occur via three primary mechanisms: incompatibility, pharmacokinetics (PK), or pharmacodynamics (PD) [1].

Incompatibility involves physical or chemical interactions affecting drug stability or bioavailability—e.g., tetracycline binds with calcium in dairy, reducing absorption [2]. PK interactions affect absorption, metabolism, distribution, or excretion; for example, grapefruit juice inhibits CYP3A4, increasing blood levels and toxicity risks of statins [3]. PD interactions modify drug effects at target receptors, enhancing or counteracting intended outcomes, as seen with vitamin K-rich foods diminishing warfarin’s anticoagulant effect [4]. Understanding and managing DFIs is thus essential for optimizing drug therapy and ensuring patient safety.

Identifying novel DFIs is critical not only clinically but also during drug development, where early detection can improve safety profiles and accelerate approval. However, experimental identification remains difficult due to the vast number of possible drug-food pairs and complex biochemical mechanisms [5]. Therefore, computational models have emerged as scalable alternatives. DFinder, for instance, uses a graph-based method integrating compound features and topological information from a large heterogeneous graph of drugs and food [5]. DFI-MS introduces a multilevel feature optimization and contrastive learning framework [6]. While promising, both models struggle with generalization to unseen drugs or food compounds. DFinder’s reliance on a fixed interaction network and DFI-MS’s a compound-wise embedding layer with a fixed set of representations hinder their ability to predict interactions for novel inputs—a critical challenge for real-world applications, particularly with new therapeutics. These limitations underscore the need for more flexible, generalizable representation approaches.

Among DFI mechanisms, cytochrome P450 (CYP450)-mediated interactions are especially important, as these enzymes metabolize 75–80% of clinical drugs [7], and account for 60–70% of DFIs [8, 9]. For example, grapefruit juice inhibits CYP3A4, increasing plasma levels of statins and antidepressants [10], while St. John’s Wort induces it, reducing efficacy of medications such as contraceptives and immunosuppressants [11]. These examples highlight the importance of incorporating CYP450 mechanisms when studying DFIs [12].

Despite CYP450’s significance, no prior computational models have explicitly used CYP450-related information for DFI prediction. This gap stems from sparse annotations for compound-CYP450 interactions. While databases such as DrugBank [13] and the Flockhart Table [14] provide some data on drug-CYP450 interactions (e.g., substrates, inhibitors), coverage remains limited, particularly for food compounds. Consequently, incorporating this critical information into existing models has been challenging. Developing sparsity-resilient models that leverage known CYP450 annotations while uncovering novel interactions is thus an important research direction.

Cross-attention mechanisms are popular in deep learning for modeling inter-modal relationships and enhancing interpretability through attention scores [15]. These scores, based on embedding similarity between *query* and *key* elements, highlight important interactions. However, attention scores can be ambiguous when no relevant *key* exists to a given *query* element, undermining interpretability [16]. ArkDTA, a drug-target interaction model, mitigates this by using attention regularization: introducing a surrogate *pseudo*-embedding that absorbs attention when no explicit relationship exists in labeled data [17].

We extend this approach to DFI prediction using CYP450 information. Here, ground-truth labels (compound-CYP450 interactions) guide attention regularization. The *pseudo*-embedding absorbs attention if a compound lacks known interaction with the CYP450 isoenzymes, improving both prediction and interpretability. Unlike ArkDTA, however, our study faces a highly sparse set of ground truth labels for CYP450 interactions, particularly for food compounds. To address this, we apply semi-supervised learning, using attention regularization only where DCI annotations exist and defaulting to standard unsupervised cross-attention otherwise. As a result, our model can predict DFIs for novel compounds while attributing interactions to specific CYP450 isoenzymes.

In this paper, we present **A**ttention **R**egularization guided by **C**YP450 Interactions for predicting **D**rug-**F**ood **I**nteractions (ArcDFI), a novel deep-learning framework that robustly predicts DFIs and enhances interpretability via CYP450-based attention mechanisms (Fig 1). Our key contributions are:

**Fig 1.**
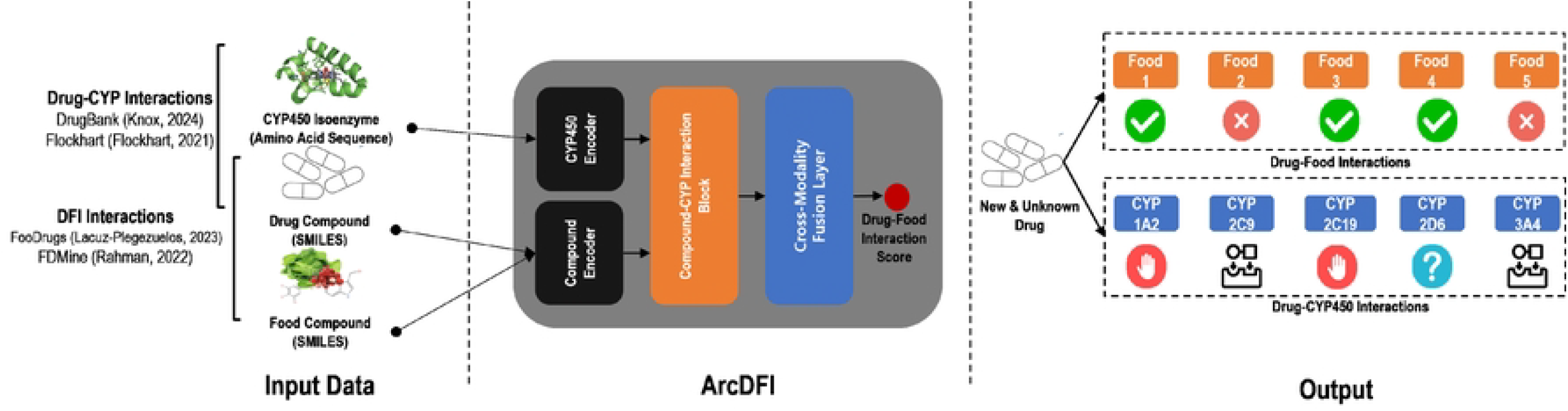
Overview of ArcDFI.

- Inspired by ArkDTA [17], we propose ArcDFI that employs attention regularization, which modulates its attention mechanism based on available information regarding drug-CYP450 interactions (DCIs).
- We demonstrate ArcDFI’s generalizability in predicting DFI interactions given unseen drugs or food compounds based on its experimental results.
- Utilizing the attention mechanism, we demonstrate ArcDFI’s explainability by finding which compound substructures are relevant in predicting DFIS and implicating which CYP450 isoenzymes are likely to form DCIs with the compound.

## Materials and Methods

### DFI Dataset Construction

The main data sources for constructing the DFI dataset were FooDrugs [18] and FDMine [19]. The latest version of the FooDrugs dataset (v4) contains over 500,000 DFIs collected from textual documents using natural language processing techniques, and inferred from gene expression data using similarity profile analysis. The FDMine dataset contains binary-labeled pairwise interactions for a unique number of 787 drugs and 563 food compounds. It is a comprehensive dataset built from two large-scale data sources: DrugBank [13] and Food Database (FooDB).

After eliminating DFIs that contain invalid compounds, we integrated both data sources to construct a large-scale DFI prediction dataset denoted as ArcDFI dataset. Positive labels indicate that a drug-food compound pair forms a metabolism-related interaction, while negative labels indicate its absence. Table 1 shows the overall statistics for the three datasets including ours.

**Table 1.** Detailed statistics for each dataset where our newly constructed ArcDFI dataset is an integration of FooDrugs and FDMine.

| Dataset | # of Drug Comp. | # of Food Comp. | # of DFI Pairs | # of Pos. Pairs | # of Neg. Pairs |
| --- | --- | --- | --- | --- | --- |
| **FooDrugs (v4)** | 17,541 | 7,676 | 166,662 | 166,662 | 0 |
| **FDMine** | 786 | 563 | 442,518 | 2,621 | 439,897 |
| **ArcDFI (ours)** | 18,325 | 8,328 | 609,180 | 169,283 | 439,897 |

### Drug-CYP450 Interaction Label Annotation

Our approach centralizes the modeling of CYP450-mediated DFIs, which requires CYP450 interaction features associated with both drug and food compounds. To augment our DFI dataset with CYP450-related information, we first gathered drug compounds annotated with CYP450 interaction labels from three data sources. The DrugBank database [13] and Drug Interactions Flockhart Table [14] provide the drug-CYP450 interactions (DCI) involving CYP450 isoenzymes and their drug interaction types. In addition, [20] released a CYP450 interaction dataset that contains both positive (presence of DCI) and negative (absence of DCI) labels for each interaction type.

After integrating the three data sources, we constructed a drug-CYP450 interaction (DCI) dataset involving five CYP450 isoenzymes (CYP1A2, CYP2C9, CYP2C19, CYP2D6, and CYP3A4), and two interaction types (substrate and inhibition). While there are other CYP450 isoenzymes (i.e., CYP2E1) and interaction types (i.e., induction), we only selected those that had relatively higher availability. The DCI dataset was used to annotate the drug compounds contained in the DFI dataset using their corresponding CYP450 interaction labels. Table 2 shows the number of drug compounds annotated for each CYP450 isoenzyme interaction type.

**Table 2.** Number of drug compounds annotated with each type of CYP450 isoenyzme (1A2, 3A4, 2C19, 2C9, 2D6) interaction (substrate, inhibition) label.

| CYP450 Isoenzyme | 1A2 |  | 3A4 |  | 2C19 |  | 2C9 |  | 2D6 |  |
| --- | --- | --- | --- | --- | --- | --- | --- | --- | --- | --- |
| Interaction Type | Sub | Inh | Sub | Inh | Sub | Inh | Sub | Inh | Sub | Inh |
| FooDrugs (v4) | 81 | 140 | 375 | 107 | 33 | 78 | 90 | 117 | 139 | 120 |
| FDMine | 92 | 398 | 329 | 132 | 57 | 167 | 78 | 209 | 170 | 212 |
| ArcDFI (ours) | 173 | 538 | 704 | 239 | 90 | 245 | 168 | 326 | 309 | 332 |

### Model Architecture for ArcDFI

As shown in Fig 2-**a)**, ArcDFI comprises the following components: Compound Substructure Encoder, Compound Graph Encoder, CYP450 Encoder, Compound-CYP Interaction Block, Cross-Modality Fusion Layer and Drug-Food Interaction Prediction Layer. It predicts a drug-food interaction score given both SMILES-represented compounds as input. Throughout this paper, the term “compound” is collectively used to refer to both drug compounds and food compounds.

**Fig 2.**
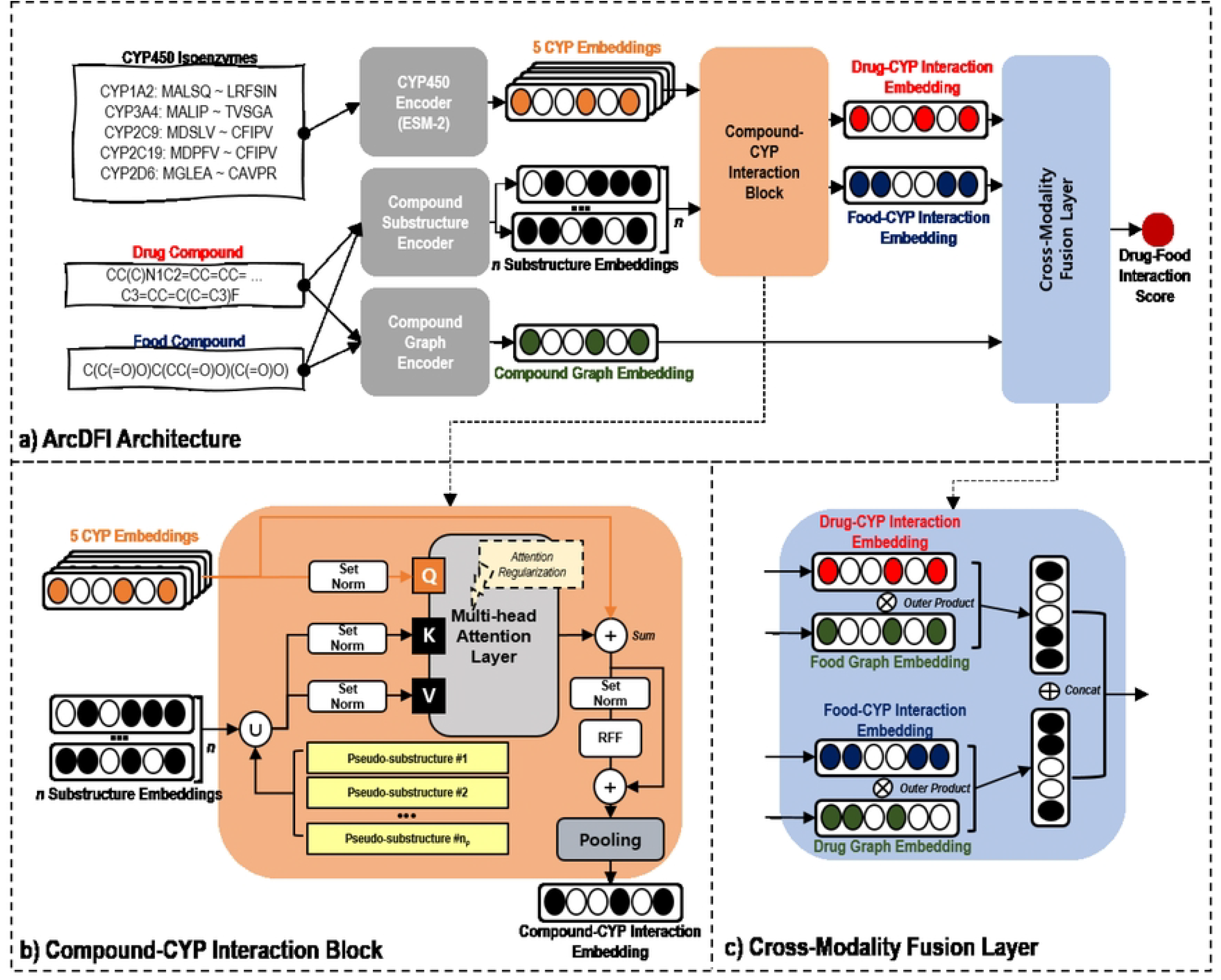
Descriptive illustration of ArcDFI. **a)** Model architecture for ArcDFI. The parameters in Compound Substructure Encoder, Compound Graph Encoder and Compound-CYP Interaction Block are shared by both drug and food compounds. **b)** Detailed illustration of the Compound-CYP Interaction Block. **c)** Detailed illustration of the Cross-Modality Fusion Layer. The drug (food) compound-CYP interaction embedding is combined with the food (drug) compound graph embedding using a vector-wise outer product, followed by concatenation of the two embeddings.

#### List of Mathematical Notations

For better understanding, we provide a list of mathematical notations base below.

|  |  |
| --- | --- |
| $d$ | Drug Compound |
| $f$ | Food Compound |
| $\mathcal{S}$ | Chemical Substructures of a Compound |
| $\mathcal{G}$ | Molecular Graph Representation of a Compound |
| $\mathcal{E}$ | CYP450 IsoEnzymes |
| $\rho$ | Trainable Pseudo-substructures |
| $\mathbf{X}^{\mathcal{S}}$ | Initial Set of $\mathcal{S}$ Embeddings |
| $\mathbf{X}^{\mathcal{G}}$ | Initial Compound Graph Representation |
| $\mathbf{X}^{\mathcal{E}}$ | Set of 5 Amino-Acid Sequences of $\mathcal{E}$ |
| $\mathbf{H}^{\mathcal{S}}$ | Set of Encoded Chemical Substructure Embeddings for a Compound |
| $\mathbf{H}^{\mathcal{G}}$ | Molecular Graph Embedding for a Compound |
| $\mathbf{H}^{\mathcal{E}}$ | Set of 5 Refined CYP450 Isoenzyme Embeddings |
| $\mathbf{H}^{\rho}$ | Set of Pseudo-Substructure Embeddings |
| $\mathbf{H}_d^{\mathcal{I}}$ | Drug Compound-Integrated CYP450 Embedding |
| $\mathbf{H}_f^{\mathcal{I}}$ | Food Compound-Integrated CYP450 Embedding |
| $\mathbf{H}^{\mathcal{F}}$ | CYP450-mediated Drug-Food Fusion Embedding |
| $\hat{\mathbf{Y}}$ | Batch of DFI Score Predictions |
| $\mathbf{Y}$ | Batch of Ground Truth DFI Labels |
| $\mathcal{A}_{i,j,k}$ | Attention Weight Score calculated between<br>the $j$th CYP450 Isoenzyme<br>and $k$th Compound Substructure<br>in the $i$th Attention Head |
| $\hat{\mathbf{Y}}_c$ | Batch of CYP450 Interaction Score Predictions |
| $\mathbf{Y}$ | Batch of Ground Truth CYP450 Interaction Labels |

#### Compound Substructure Encoder

The Compound Substructure Encoder first takes a SMILES-represented compound as input and converts it into 1024-dimensional Extended Connectivity Fingerprints (ECFPs) with a radius set to 2. Because each bit position of its binary ECFP represents the presence of its substructure detected within a certain radius, we utilized this information by converting each compound into a set of 2*h*-dimensional trainable substructure embeddings [17]. These initial substructure embeddings corresponding to the drug and food compounds were then propagated through an MLP, resulting in a set of encoded *n*_*d*_ and *n*_*f*_ chemical substructure embeddings, respectively, where *d* and *f* represent a drug compound and a food compound respectively 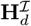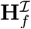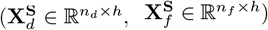.

The Compound Substructure Encoder that takes the ECFPs from SMILES representation of drugs and food compounds as inputs is mathematically expressed as follows:

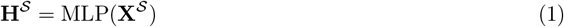

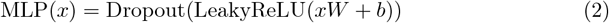

where **X**^*𝒮*^ ∈ ℝ ^*n×*2*h*^ is an initial set of 2*h*-dimensional chemical substructure embeddings converted from the compound’s ECFP representation. MLP comprises a linear transformation layer (*W* ∈ ℝ^2*h×h*^, *b* ∈ ℝ^*h*^) followed by non-linear activation function LeakyReLU and Dropout layer whose dropout rate is globally set to 0.3.

#### Compound Graph Encoder

The Compound Graph Encoder also uses a SMILES-represented compound as input and converts it into its molecular graph representation (**X**^*𝒢*^) compatible with its inherent Graph Isomorphism Network (GIN) convolution layer that additionally incorporates edge attributes [21]. The initial features for the nodes (atoms) are the atomic number, chirality, degree, formal charge, number of hydrogen atoms, number of radical electrons, hybridization, aromaticity, and ring-like structure. The initial features of the edges (bonds) are the bond types, stereo configuration, and conjugation.

Atom-wise node embeddings of the given compound were built using the GIN convolution layer based on the initial atom-bond information and topological characteristics. They are subsequently aggregated through a global mean pooling layer into an *h*-dimensional single-handed molecular graph embedding which is mathematically expressed as 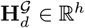 and 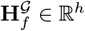 respectively.

The Compound Graph Encoder, which uses the SMILES representation of the drug and food compounds as input is mathematically expressed as follows:

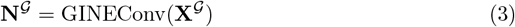

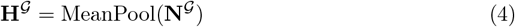

where **X**^*𝒢*^ is the molecular graph of the input compound previously converted from its SMILES representation and contains the initial atom and bond features. GINEConv is the GIN convolution layer using edge attribute features, and MeanPool is a global mean pooling layer that aggregates the node embeddings **N**^*𝒢*^ to a single molecular graph embedding **H**^*𝒢*^ ∈ ℝ^*h*^.

#### CYP450 Encoder

The CYP450 Encoder takes a set of five CYP450 isoenzymes initially represented as amino acid sequences as inputs and converts them into a set of contextualized CYP450 isoenzyme embeddings. The input amino acid sequence representations for the isoenzymes were obtained from the UniProt database [22]. They were fed into ESM-2, a large-scale protein language transformer model trained on millions of protein sequences [23], which resulted in a set of five 480-dimensional language-based embeddings **L**^*ε*^ ∈ ℝ ^5*×*480^. Lastly, these language-based embeddings are propagated through an MLP, which results in a set of 5 *h*-dimensional refined protein embeddings **H**^*ε*^ ∈ ℝ ^5*×h*^. Note that the contextualized CYP450 isoenzyme embeddings are universally used for all drug-food compound pairs within the same batch during training.

The CYP450 Encoder, which takes the five CYP450 isoenzymes **X**^*ε*^, each represented as an input sequence of amino acids, is mathematically expressed as follows:

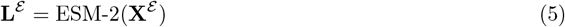

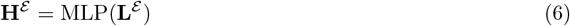

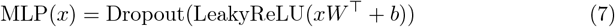

where **L**^*ε*^ ∈ ℝ ^5*×*480^ are the language-based embeddings for the five CYP450 isoenyzmes encoded by the protein language transformer model, ESM-2 (esm2-t12-35M-UR50D). MLP comprises a linear transformation layer (*W* ∈ ℝ ^480*×h*^, *b* ∈ ℝ^*h*^) followed by LeakyReLU and a Dropout layer.

#### Compound-CYP Interaction Block

The Compound-CYP Interaction Block (Fig 2-**b)**) calculates multi-head cross-attention scores between the contextualized CYP450 isoenzyme and chemical substructure embeddings, which are the input queries and key values, respectively. Its design intuition aligns with a ligand (food or drug compound) binding to the target CYP450 isoenzyme, exhibiting either inhibitory or substrate-related effects. By interpreting pairwise attention scores calculated by the cross-attention mechanism, we can imply which substructures play a crucial role in DCIs.

The cross-attention mechanism usually involves the calculation of pairwise attention weights between CYP450 isoenzymes (queries) and chemical substructure embeddings (keys) in an unsupervised manner. Attention weights are distributed to the keys for each query, regardless of whether the compound actually interacts with the isoenzyme. To address this issue, we adopted ArkDTA’s attention regularization method, which aims to modulate the distribution of attention weights driven by an auxiliary loss objective [17].

Specifically, a set of *n*_*ρ*_ trainable *h*-dimensional pseudo-substructure embeddings are first appended to the current set of chemical substructure embeddings. Attention weights distributed to the pseudo-substructures imply absence of DCIs while the opposite applies for those distributed to the actual ones. We treated the sum of the weights assigned to both the pseudo-substructures and the actual substructures as binary class probability scores. They are fed to a binary cross-entropy loss function, where the ground truth labels are the compound-CYP450 interaction annotations from our DCI dataset.

Given that our study included two types of DCI substrates and inhibition, we applied supervised attention regularization to the attention weights from the first and second heads of the block, corresponding to substrate and inhibition effects, respectively. The remaining attention heads were unsupervised during the training process. The output of the Compound-CYP Interaction Block is an aggregation of 5 *h*-dimensional attention-based CYP450 isoenzyme embeddings **O**^*ε*^ ∈ ℝ^5*×h*^ using mean pooling. We denote this final output as compound-integrated CYP450 embedding **H**^*ℐ*^ ∈ ℝ^*h*^. A detailed explanation of the loss objective is available in the Auxiliary Loss Objective section.

The Compound-CYP Interaction Block that takes the set of five contextualized CYP450 isoenzymes **H**^*ε*^ and *n* chemical substructure embeddings **H**^*𝒮*^ as inputs is mathematically expressed as follows:

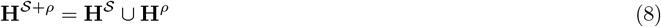

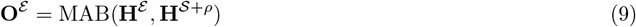

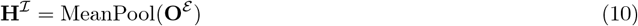

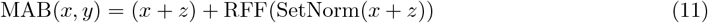

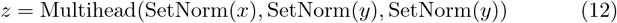

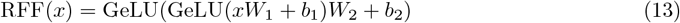

where MAB is a multihead attention block borrowed from the Set Transformer architecture [24] enhanced by set normalization layers and equivariant skip connections [25]. The attention block includes a multihead attention layer, where the input query, key, and value embeddings are **H**^*ε*^, **H**^*𝒮* +*ρ*^ and **H**^*S*+*ρ*^ respectively. It employs four attention heads with pairwise attention weights computed using additive attention. Note that the computed attention weights from the first two heads are fed to an auxiliary loss objective. RFF is a row-wise feedforward layer consisting of two layers of linear transformation (*W*_1_, *W*_2_ ∈ ℝ^*h×h*^, *b*_1_, *b*_2_ ∈ ℝ^*h*^) each followed by Gaussian Error Linear Unit (GeLU) activation function.

#### Cross-Modality Fusion Layer

To effectively model the complex interactions between CYP450 isoenzymes, drug compounds, and food compounds, we implemented a Cross-Modality Fusion Layer (Fig 2-**c)**) that combines attention-based drug-CYP450 integrative features 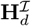 with the food compound’s graph topological features 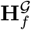 and vice versa 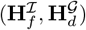.

Recall that 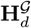 and 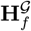 represent molecular graph embeddings for the drug and food compound, respectively, whereas 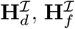 denote the CYP450 embeddings integrated with the drug and food compound, respectively.

The fusion process employs a vector-wise outer product between the two heterogeneous embeddings for both sides of the drug-food pair. Each of the outer product results is reshaped into a single *h*^2^-dimensional embedding. Lastly, the two reshaped embeddings are concatenated to each other vector-wise, producing a final output of this layer: a 2*h*^2^-dimensional CYP450-mediated drug-food fusion embedding 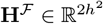.

The Cross-Modality Fusion Layer that takes the compound-integrated CYP450 isoenzyme embeddings 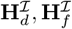 and molecular graph embeddings 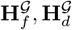 as input is mathematically expressed as follows,

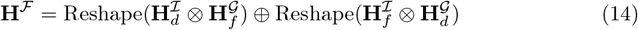

where Reshape, ⊗,⊕ refers to the reshaping process from ℝ^*h×h*^ matrix to 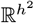 vector, vector-wise outer product and concatenation.

#### Drug-Food Interaction Prediction Layer

Finally, the Drug-Food Interaction Prediction Layer takes the single CYP450-mediated drug-food fusion embedding as input and ultimately predicts the interaction likelihood between the drug and food compound. The Drug-Food Interaction Prediction Layer is a deeply stacked feedforward layer that takes the single CYP450-mediated drug-food fusion embedding as input and ultimately predicts the interaction likelihood between the drug and food compound. This can be mathematically expressed as follows:

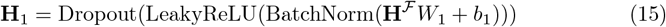

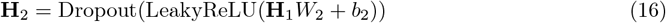

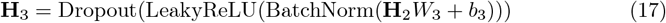

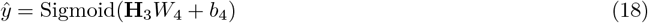

where BatchNorm and Sigmoid refer to the batch normalization and sigmoid functions, respectively. Weights and bias for the linear transformation layers are 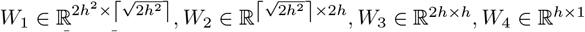 and 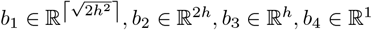 respectively.

### Model Optimization

ArcDFI is trained under two loss objectives which are the primary loss objective for predicting DFIs and auxiliary loss objective for attention regularization. The former is implemented under supervised learning setting,

#### Primary Loss Objective

The batch-wise primary loss objective for drug-food interaction prediction, treated as a binary classification task, is mathematically expressed as follows:

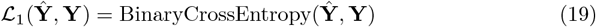

where **Ŷ** ∈ ℝ^*b*^ and **Y** ∈ {0, 1} ^*b*^ is a batch of predicted DFI scores and ground truth binary labels respectively, and *b* is the batch size.

#### Auxiliary Loss Objective

To enhance ArcDFI’s understanding of CYP450-mediated drug-food interactions through semi-supervised learning, we introduced an auxiliary loss objective to regularize the attention mechanism. Let 𝔸_*i,j,k*_ denote the attention weights computed between the *j*th CYP450 isoenzyme and *k*th compound substructure in the *i*th attention head within the Compound-CYP Interaction Block. The value of 𝔸_*i,j,k*_ can be interpreted as the likelihood of the *k*th compound substructure contributing to the substrate (*i* = 1) or inhibitory effects (*i* = 2) to the *j*th CYP450 isoenzyme. To clarify, we applied attention regularization to the first (𝔸_1,*j,k*_) and second attention heads (𝔸_2,*j,k*_), whereas the remainder were left unsupervised as originally designed.

As described earlier, the Compound-CYP Interaction Block module appends a universal set of *n*_*ρ*_ trainable pseudo-substructure embeddings to the current set of *n* key substructure embeddings resulting in a total of *n* + *n*_*ρ*_ embeddings involved in pairwise attention weight computation with respect to the five query CYP450 isoenzyme embeddings. Given 𝔸_1,*j,k*_, if the partner compound forms a substrate binding interaction with the *j*th CYP450 isoenzyme according to the ground truth label in our DCI dataset, our designed auxiliary loss objective encourages the distribution of attention weights to concentrate on the *n* actual substructure embeddings 𝔸_1,*j*,1:*n*_, while reducing the weights for the *n*_*ρ*_ pseudo-substructure embeddings, 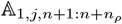. Similarly, this modulation applies to 𝔸 for compounds exhibiting inhibitory effects on the *j*th CYP450 isoenzyme.

Conversely, when the ground truth label for a specific DCI type is negative, the loss objective promotes the opposite behavior by diverting attention away from the actual substructure embeddings. If the DCI label is unavailable, the loss objective is not applied, leaving the attention distribution unsupervised. Note that the calculated pairwise attention weights for each row in 𝔸_*i,j,k*_ are normalized using a softmax function, making them inherently suitable for optimization using a cross-entropy loss in the auxiliary objective.

The batch-wise auxiliary loss objective for attention regularization based on compound-CYP450 interactions is mathematically expressed as follows:

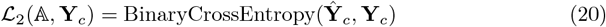

where **Ŷ** _*c*_ ∈ ℝ^*b×*10^ and **Y**_*c*_ ∈ {0, 1}^*b×*10^ is a batch of predicted interaction scores and ground truth binary labels respectively for each CYP450 isoenzyme and type. The interaction scores **Ŷ**_*c*_ are calculated based on the computed attention weights extracted from the Compound-CYP Interaction Block.

Note that the sparse proportion of drug compounds with CYP450 isoenzyme interaction labels sets the auxiliary loss objective in a semi-supervised learning setting. When interaction labels are unavailable, the corresponding predicted interaction scores remain unsupervised. Only the drug compounds with available interaction labels are used to update the parameters of the Compound-CYP Interaction Block. Fig 3 provides an illustrated description for the auxiliary loss objective.

**Fig 3.**
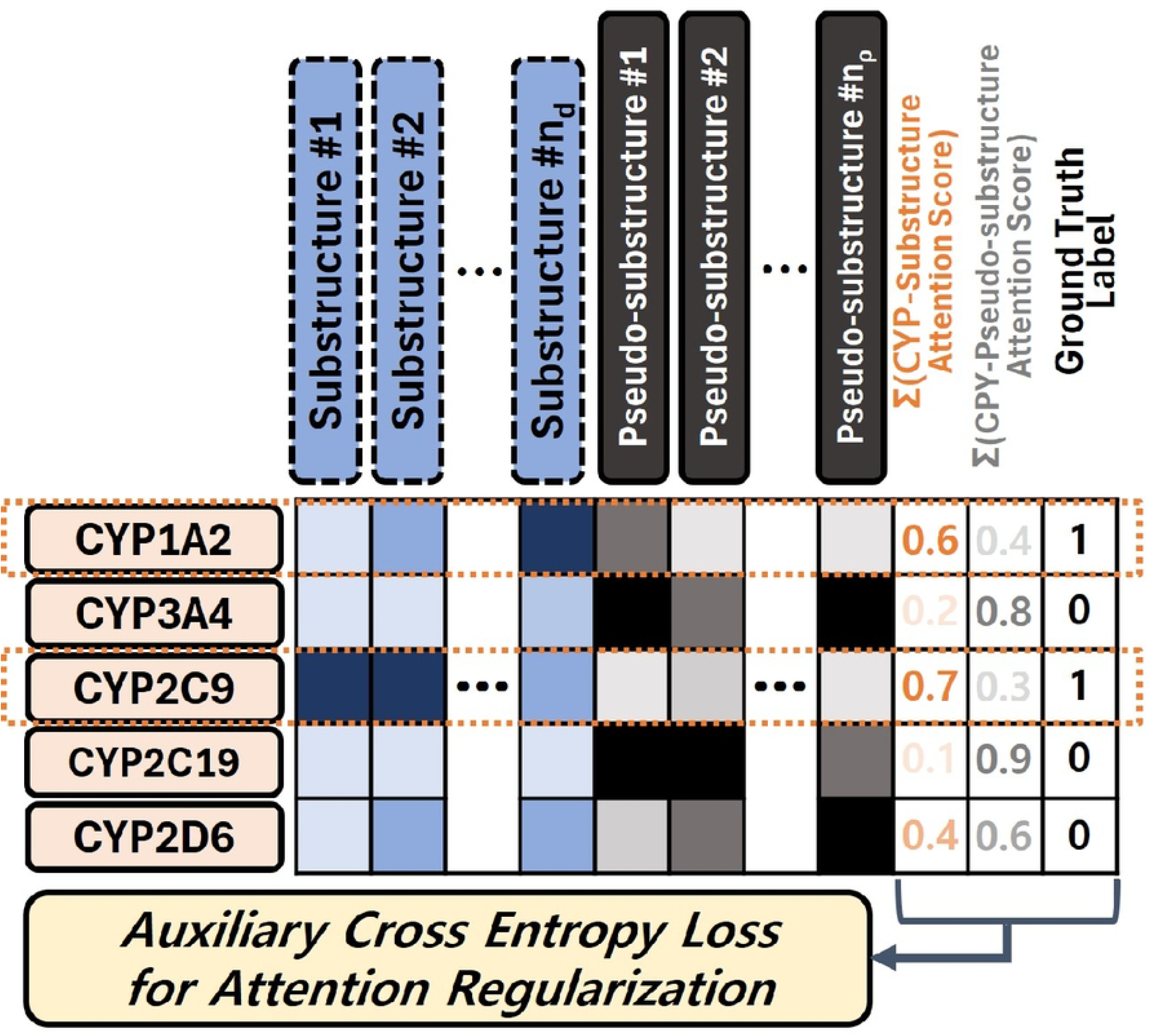
Detailed illustration for Attention Regularization Auxiliary Loss Objective.

#### Training ArcDFI

Conclusively, the batch-wise total loss objective for training ArcDFI is mathematically expressed as follows,

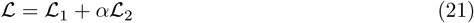

where *α* is the auxiliary loss coefficient for controlling the intensity of attention regularization based on CYP450 interaction.

We trained ArcDFI for a maximum of 100 epochs using early stopping, optimized with the AdamW optimizer. The batch size (*b*), learning rate, and weight decay were set to 1024, 0.0001, and 0.0001, respectively. Early stopping was configured with a patience of 10 epochs, using the validation loss as the monitoring metric. The auxiliary loss coefficient *α* was set to 5. All hyperparameters for the model architecture and training algorithm were consistently the same across all settings involving ArcDFI. For the model-related configurations of ArcDFI, the hidden dimensional size of embeddings and the number of pseudo-substructure embeddings were set to *h* = 128 and *ρ* = 10.

## Results

### Evaluation on DFI Prediction

To evaluate ArcDFI’s performance in DFI prediction, we conducted experiments using 10 baseline models which are DeepSynergy [26], DeepDDI [27], EPGCN-DS [28], CASTER [29], SSI-DDI [30], MatchMaker [31], MR-GNN [32], DeepDrug [27], DFinder [5] and DFI-MS [6]. While only DFinder and DFI-MS were developed with the sole purpose of predicting DFIs, we included models originally designed for drug-drug interaction prediction in this experiment, as described by [6].

Unlike previous studies on DFI, we employed a stricter evaluation approach that involved splitting the dataset based on its compound clusters. We first utilized Butina clustering to build drug and food compound clusters in the DFI dataset [33] and then made two data splits based on the food or drug compound clusters, which are referred to as cold drug and cold food settings. This data split approach has been widely used in research on drug-target interaction prediction.

We repeated the experiments three times using different random seeds for each model and data-split setting to ensure stability and robustness. We used five evaluation metrics to measure the predictive performance: Area Under ROC Curve (**AUROC**), Area Under Precision-Recall Curve (**AUPRC**), F1-score (**F1**), Precision, and Recall. The final scores for each model were averaged across three runs for each split setting.

Tables 3 and 4 show the quantitative results of evaluating the new drug and food setting, respectively. ArcDFI outperformed its baseline models for the cold drug setting, especially in terms of AUROC, AUPRC, and F1, and even its ablated version. While cold food evaluation setting presented a harder challenge to the DFI models, ArcDFIshowed second-best predictive performance. In contrast, the ablated version of ArcDFI, in which its auxiliary loss objective was removed, outperformed other models in the cold food setting. These results demonstrated that the proposed attention regularization method is advantageous for improving the generalizability of unseen drug compounds. In contrast, as our dataset does not contain any CYP information associated with food compounds, ArcDFI has limited generalizability in unseen foods because its Compound-CYP Interaction Block relies solely on traditional unsupervised cross-attention mechanisms.

**Table 3.** Evaluation results for ArcDFI, its ablated version, and baseline models under the cold drug experiment setting. All evaluation scores were averaged over three iterations along with their standard deviation. Best results are bold-faced.

| Model | Cold Drug Evaluation Setting |  |  |  |  |
| --- | --- | --- | --- | --- | --- |
|  | AUROC | AUPRC | F1 | Precision | Recall |
| DeepSynergy | 0.8638±0.0006 | 0.8122±0.0047 | 0.6591±0.0061 | 0.5155±0.0080 | 0.9138±0.0027 |
| DeepDDI | 0.9048±0.0350 | 0.9151±0.0511 | 0.7982±0.0398 | 0.7078±0.0532 | 0.9151±0.0196 |
| EPGCN-DS | 0.9050±0.0001 | 0.8454±0.0003 | 0.7125±0.0057 | 0.8411±0.0046 | 0.6181±0.0111 |
| CASTER | 0.7660±0.0058 | 0.6478±0.0083 | 0.5861±0.0158 | 0.4381±0.0232 | 0.8872±0.0226 |
| SSI-DDI | 0.9697±0.0058 | 0.9451±0.0124 | 0.8816±0.0155 | 0.8884±0.0096 | 0.8757±0.0370 |
| MatchMaker | 0.8883±0.0390 | 0.8472±0.0704 | 0.6685±0.0487 | 0.5258±0.0636 | 0.9237±0.0065 |
| MR-GNN | 0.9674±0.0017 | 0.9388±0.0022 | 0.8733±0.0092 | 0.8936±0.0139 | 0.8546±0.0292 |
| DeepDrug | 0.9615±0.0021 | 0.9364±0.0066 | 0.8551±0.0130 | **0.9053±0.0167** | 0.8111±0.0341 |
| DFinder | 0.5002±0.0094 | 0.0011±0.0001 | 0.0022±0.0001 | 0.0011±0.0001 | 0.9446±0.0682 |
| DFI-MS | 0.7095±0.0111 | 0.5614±0.0097 | 0.5543±0.0023 | 0.3917±0.0011 | **0.9480±0.0065** |
| ArcDFI ($-\mathcal{L}_2$) | 0.9474±0.0134 | 0.8762±0.0517 | 0.8385±0.0153 | 0.7775±0.0285 | 0.9104±0.0104 |
| ArcDFI (ours) | **0.9717±0.0034** | **0.9494±0.0082** | **0.8833±0.0056** | 0.8761±0.0196 | 0.8911±0.0165 |

**Table 4.** Evaluation results for ArcDFI, its ablated version and baseline models under the cold food experiment setting. All evaluation scores were averaged over three iterations along with their standard deviation. Best results are bold-faced.

| Model | Cold Food Evaluation Setting |  |  |  |  |
| --- | --- | --- | --- | --- | --- |
|  | AUROC | AUPRC | F1 | Precision | Recall |
| DeepSynergy | 0.8013±0.0047 | 0.6070±0.0159 | 0.5956±0.0055 | 0.4612±0.0078 | 0.8405±0.0040 |
| DeepDDI | 0.7624±0.0102 | 0.4940±0.0195 | 0.6074±0.0081 | 0.4697±0.0105 | 0.8596±0.0176 |
| EPGCN-DS | 0.8665±0.0021 | 0.7031±0.0154 | 0.6687±0.0054 | 0.6985±0.0039 | 0.6415±0.0131 |
| CASTER | 0.8961±0.0074 | 0.7974±0.0319 | 0.6958±0.0064 | 0.5798±0.0128 | 0.8701±0.0099 |
| SSI-DDI | 0.9311±0.0071 | 0.8442±0.0291 | 0.7938±0.0086 | 0.7896±0.0087 | 0.7986±0.0253 |
| MatchMaker | 0.7620±0.0516 | 0.5829±0.1023 | 0.5395±0.0135 | 0.3962±0.0056 | 0.8459±0.0415 |
| MR-GNN | 0.9219±0.0021 | 0.8046±0.0140 | 0.7791±0.0134 | 0.7859±0.0383 | 0.7766±0.0599 |
| DeepDrug | 0.9366±0.0038 | 0.8565±0.0176 | 0.7821±0.0107 | **0.8574±0.0083** | 0.7191±0.0169 |
| DFinder | 0.2870±0.0072 | 0.0003±0.0000 | 0.0010±0.0000 | 0.0005±0.0000 | **0.9989±0.0006** |
| DFI-MS | 0.6061±0.0280 | 0.3905±0.0405 | 0.4680±0.0050 | 0.3138±0.0041 | 0.9202±0.0095 |
| ArcDFI ($-\mathcal{L}_2$) | **0.9462±0.0069** | **0.8586±0.0155** | **0.8129±0.0094** | 0.7652±0.0086 | 0.8673±0.0207 |
| ArcDFI (ours) | 0.9321±0.0073 | 0.8074±0.0362 | 0.8029±0.0136 | 0.7648±0.0442 | 0.8471±0.0231 |

Moreover, our model design approach, which features protein language modeling for CYP450 isoenzymes, the mixture of two compound modalities (molecular graph, substructures) and efficient parameter sharing between two compound inputs (Chemical Substructure Embedding Layer, Compound Graph Encoder, CYP Interaction Block), proved to be the optimal choice for this DFI prediction task. Whereas the other baseline models did not utilize CYP450 information or cross-modality fusion, our model architecture exhibited superiority in both cold drug and food evaluation settings.

Although DFinder and DFI-MS were expected to perform better than the DDI prediction models, they exhibited poor performance when evaluated in the new drug and food setting. The poor performances of both models are related to their inherent model structures. DFinder is a graph network-based model that relies on network topology for learning compound features represented as node embeddings. Similarly, DFI-MS depends on a set of embeddings that are only partially updated during training, leaving the model unable to generalize well to novel compounds, resulting in reduced performance in the evaluated settings. We remark that these two models require to be retrained on newly introduced compounds but ours can be directly utilized by running inference on such cases.

### Analysis on Attention Weights

To investigate the underlying relationships between CYP450 isoenzymes and drug-food compounds, we performed model inference and visualized the calculated attention weights extracted from ArcDFI’s Compound-CYP Interaction Block for each compound (drug, food) and head (substrate, inhibition) using heatmaps, as shown in Fig 4**-(a),(b),(e),(f)** and Fig 5**-(a),(b),(e),(f)**. Red cells indicate the pairwise interaction scores between the drug compound and each CYP450 isoenzyme under a specific interaction type (substrate or inhibition). Note that the pairwise scores were calculated based on the summation of attention weights distributed to each compound substructure, with respect to each CYP450 isoenzyme. Blue cells indicate the contribution of each chemical substructure of a compound to CYP450 isoenzyme interactions. The chemical substructures in the heatmap were represented using SMILES.

**Fig 4.**
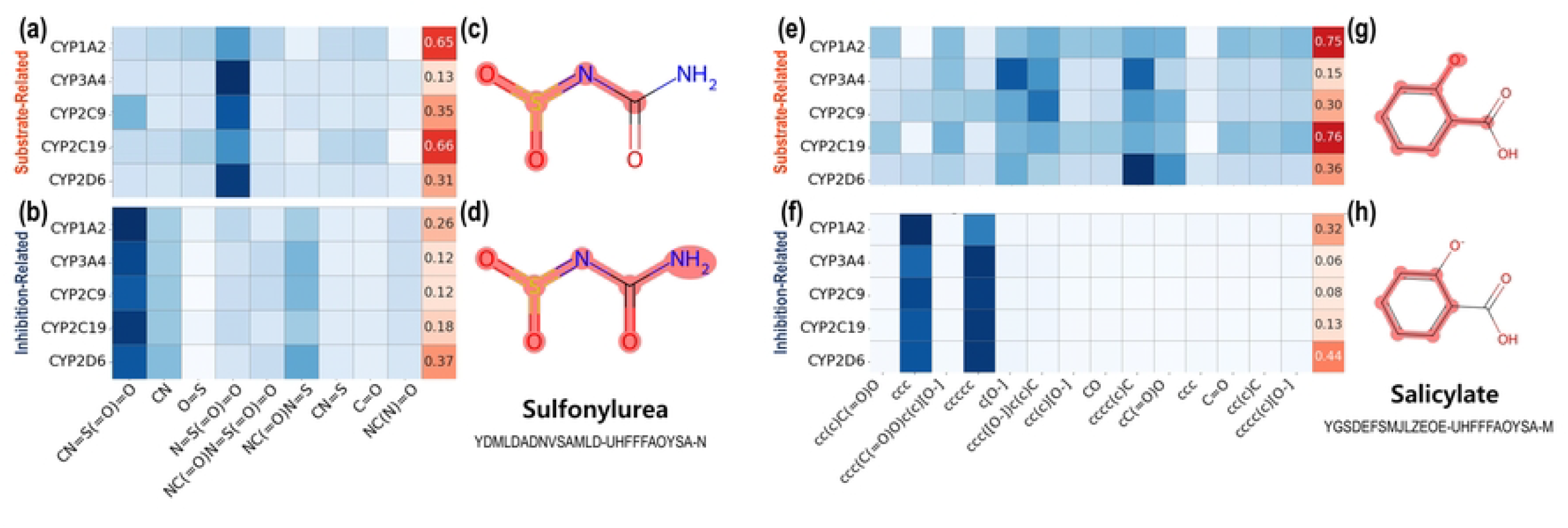
Analysis on ArcDFI’s Compound-CYP Interaction Block for drug-food compound pair Sulfonylurea and Salicylate. **(a), (b):** Attention weights and highlighted compound substructures extracted from ArcDFI’s Compound-CYP Interaction Block head related to substrate-type interactions for Sulfonylurea, respectively. **(c), (d):** Attention weights and highlighted compound substructures extracted from ArcDFI’s Compound-CYP Interaction Block head related to inhibition-type interactions for Sulfonylurea, respectively. **(e), (f):** Attention weights and highlighted compound substructures extracted from ArcDFI’s Compound-CYP Interaction Block head related to substrate-type interactions for Salicylate, respectively. **(g), (h):** Attention weights and highlighted compound substructures extracted from ArcDFI’s Compound-CYP Interaction Block head related to inhibition-type interactions for Salicylate, respectively.

**Fig 5.**
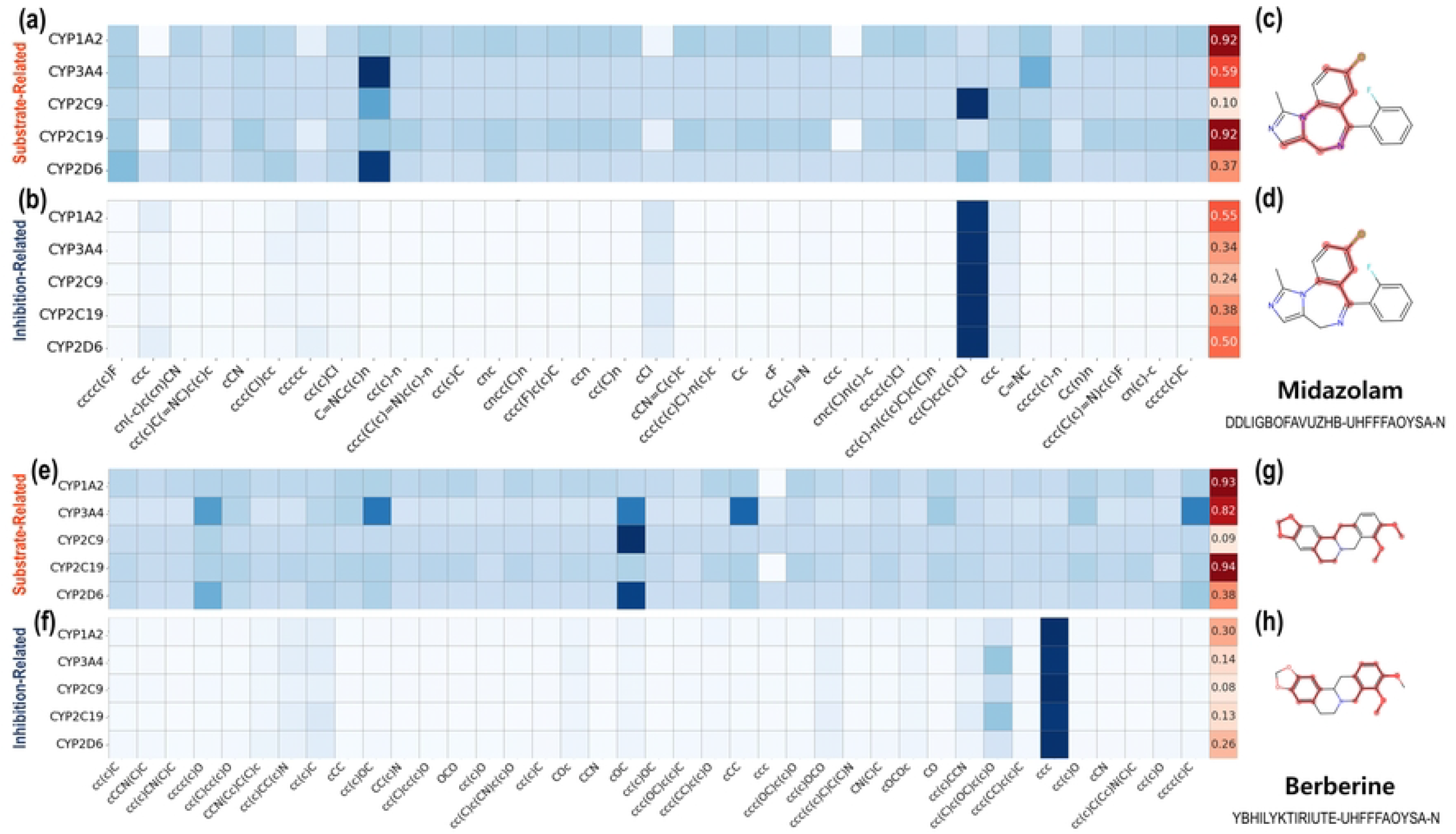
Analysis on ArcDFI’s Compound-CYP Interaction Block for drug-food compound pair Midazolam and Berberine. **(a), (b):** Attention weights and highlighted compound substructures extracted from ArcDFI’s Compound-CYP Interaction Block head related to substrate-type interactions for Midazolam, respectively. **(c), (d):** Attention weights and highlighted compound substructures extracted from ArcDFI’s Compound-CYP Interaction Block head related to inhibition-type interactions for Midazolam, respectively. **(e), (f):** Attention weights and highlighted compound substructures extracted from ArcDFI’s Compound-CYP Interaction Block head related to substrate-type interactions for Berberine, respectively. **(g), (h):** Attention weights and highlighted compound substructures extracted from ArcDFI’s Compound-CYP Interaction Block head related to inhibition-type interactions for Berberine, respectively.

Since the substructure-wise attention weights assigned to each CYP isoenzyme may not fully provide model interpretability, we visualized each compound with its important substructural regions being colored with red, as shown in Fig 4**-(c),(d),(g),(h)** and Fig 5**-(c),(d),(g),(h)**. We first extracted the top three attended substructures from our heat maps and mapped each atom onto the molecule. Finally, we highlighted the designated atoms with red color, which represent the substructural regions focused on by ArcDFI’s Compound-CYP Interaction Block.

Fig 4 displays the attention weights and highlighted substructures extracted from the ArcDFI’s Compound-CYP Interaction Block when given a pair of unseen drug compound and seen food compound, Sulfonylurea and Salicylate, as input. Sulfonylurea is a drug-like compound that has been traditionally used for treating diabetes, while Salicylate is an organic acid that is known to exhibit analgesic, antipyretic, and anti-inflammatory effects [34, 35].

According to Fig 4**-(a)**, the weights imply that Sulfonylurea forms a DCI with CYP1A2 and CYP2C19 [36] (0.65, 0.66) as a metabolized substrate while its substructure N=S(=O)=O seemed to exhibit a relatively high influence on this interaction process. On the contrary in Fig 4**-(b)**, Sulfonylurea exerted a relatively weaker inhibitory effect on CYP450 isoenzymes, with CYP2D6 being the most affected (0.37).

In Fig 4**-(e)**, the weights indicate that Salicylate forms a DCI with CYP1A2 and CYP2C19 as substrates, which can potentially have stronger implications for a DFI between two compounds sharing the same isoenzymes as substrates (0.75, 0.76). In addition, the attention weights in Fig 4**-(f)** indicate that Salicylate does not have inhibitory interactions with the CYP450 isoenzymes. Interestingly, the CYP2D6 isoenzyme received the highest score among the five isoenzymes (0.44), which was the same for Sulfonylurea.

The highlighted substructures shown in Fig 4**-(c)** and **-(d)** indicate that most of the substructural regions in Sulfonylurea are perceived to contribute to the overall attention weights, which aligns with the fact Sulfonylureas is used as a chemical derivative for making other drug compounds. As shown in Fig 4**-(g)** and **-(h)**, the aromatic substructures of Salicylate have received the most attention weights overall, which may implicate *π*-*π* interactions with the CYP450 isoenzyme targets.

Fig 5**-(a),(b),(e),(f)** shows the visualized attention weights when ArcDFI was given an out-of-dataset DFI pair (neither the drug nor the food compound was in our DFI dataset), with Midazolam and Berberine, as input. Midazolam is a sedative used for surgical purposes [37] and is widely used as a probe substrate for CYP3A4 isoenzyme [38], while Berberine is a plant-derived organic compound known to have anti-cancer effects [39].

As shown in Fig 5**-(a)** and **-(e)**, the attention weights indicated that both Midazolam and Berberine actively interacted with three CYP450 isoenzymes (1A2, 3A4, 2C19) as substrates. This aligns with our previous observation that ArcDFI recognizes a drug-food compound pair with an interaction when the two sides share the same CYP isoenzymes as metabolized substrates. Interestingly, no common molecular substructures exhibited similar DCI patterns, as shown in Fig 5**-(c)** and **-(g)**.

Interestingly, ArcDFI perceived Midazolam as a possible inhibitory agent targeting CYP1A2 and CYP2D6 (0.55, 0.50) as shown in Fig 5**-(b)**. While this may require further investigation, we can expect ArcDFI to suggest novel DCIs and DFI predictions for unexplored drug-food pairs.

In conclusion, ArcDFI can highlight the parts of a drug or food compound most likely to interact with particular CYP450 isoenzymes. By investigating the attention weights extracted from its Compound-CYP Interaction Block, we can derive various DCI-related hypotheses to explain or refine the DFI predictive framework.

## Discussion

### Limitations

One of the main limitations of this study was the constructed DFI and DCI datasets. While the DFI dataset may seem to contain a large number of DFI pairs (609,180), it still suffers from sparsity issues as the total number of possible DFI pairs is 152,610,600, resulting in a sparsity rate of 0.4%. Despite our efforts to gather all available DCIs from multiple data sources, the number of drug compounds annotated with CYP450 isoenzyme interactions remain extremely sparse. Also, other interaction types (i.e., inducing effects) and CYP450 isoenzymes (i.e., CYP2E1) should be considered. As some studies state that Salicylate induces CYP2C19 [40], we expect our model’s Compound-CYP Interaction Block to align with this perspective, only if its attention regularization is augmented by sufficient annotated DCI data related to induction-type interactions.

The absence of available CYP interaction labels for food compounds restricts the potential of the attention regularization method, as shown in the evaluation results for the cold food setting experiments. For instance, the attention weights related to the interaction-type interaction of Berberine with CYP450 isoenzymes were inconsistent with experimental studies [41, 42]. This is because the Compound-CYP Interaction Block relies only on the DCI information for modulating its inherent attention mechanism, which lacks guidance from Food-CYP450 interaction (FCI) information.

Another limitation is related to the use of compound substructures originating from the ECFPs of drug and food compounds. Although this approach is computationally efficient, the structural information represented by these substructures is not canonical owing to the inherent hashing process in ECFPs. That is, two identical embeddings may represent different molecular substructures. Additionally, the resolution of ECFPs is constrained by a radius of 2, potentially overlooking larger yet meaningful substructures. Although the compound graph encoder somewhat mitigates these potential weaknesses, there is still room for improvement in the representation of the substructures for a given compound.

The main premise of our study was the assumption that DFIs are primarily related to CYP450-mediated drug metabolism. While this assumption, implemented as an auxiliary loss objective in ArcDFI, demonstrated strong empirical performance in our cold drug experiments, many other factors influencing DFIs should be considered. Non-CYP enzymes or pathways are involved in the oxidative metabolism of food compounds [43]. Certain food products can influence microbiota, which may alter the metabolism of drug compounds and vice versa [44]. Furthermore, food can affect drug absorption by influencing the patient’s gastrointestinal physiology. This necessitates annotating not only DFI types but also their underlying mechanisms, such as CYP450 isoenzymes or other relevant factors.

### Future Work

We propose several directions for future improvements to ArcDFI. First, we plan to utilize automated text-mining approaches leveraging Large Language Models to collect more experimentally known DFIs and CYP450 interactions from the biomedical literature, and to improve the dataset quality through manual curation performed by domain experts. Furthermore, we plan to employ molecular docking simulations and binding affinity prediction tools to augment the unlabeled DCIs in our datasets. These additional data sources enhance the utility of attention regularization, thereby improving the training and predictive performance of ArcDFI.

Second, we plan to seek ways to improve the current design of the ArcDFImodel architecture, specifically related to its molecular representation learning and attention regularization method. In particular, we plan to explore better alternatives for representing compound substructures using other fingerprint-based representations, such as pharmacophores, molecular fragmentization (BRICS Decomposition [45], RECAP Algorithm [46]), or SELFIES representation [47]. Moreover, we plan to implement task-specific modifications to the attention layers and devising robust optimization strategies to maximize the synergistic effects of attention regularization and DFI prediction.

Lastly, we plan to incorporate biological experiments to verify ArcDFI’s DFI predictions and DCI hypotheses derived from its attention mechanism, when given an out-of-dataset drug-food pair as input for its model inference. This step is critical for strengthening the reliability and applicability of the model in real-world scenarios and for bridging computational predictions with clinical practice.

## Conclusion

We introduce a novel DFI prediction model ArcDFI, by incorporating attention regularization guided by compound-CYP450 interactions, which offers improved generalizability and interpretability in DFI prediction. Despite the challenges posed by dataset sparsity and the limited availability of CYP450 interaction labels, our model demonstrated promising results in terms of both predictive performance and interpretability. Also, the attention weights extracted from our model’s Compound-CYP Interaction Block provided fresh insights into novel drug-food and compound-CYP450 relationships. We expect our model to facilitate discovery of novel drug-food interactions.

